# Real-Time Structure Search and Structure Classification for AlphaFold Protein Models

**DOI:** 10.1101/2021.10.21.465371

**Authors:** Tunde Aderinwale, Vijay Bharadwaj, Charles Christoffer, Genki Terashi, Zicong Zhang, Rashidedin Jahandideh, Yuki Kagaya, Daisuke Kihara

## Abstract

AlphaFold2 showed a substantial improvement in the accuracy of protein structure prediction. Following the release of the software, whole-proteome protein structure predictions by AlphaFold2 for 21 organisms were made publicly available. Here, we developed the infrastructure, 3D-AF-Surfer, to enable real-time structure-based search for the AlphaFold2 models by combining molecular surface representation with 3D Zernike descriptors and deep neural networks.

---

Structural biology has entered a new phase when structure prediction methods, particularly a recent method, AlphaFold2^1^, consistently produce reliable computational structure models with atomic accuracy. Protein structure prediction has been extensively studied in the computational biology community. Taking advantage of the accumulated protein sequence and structure information in the Protein Data Bank (PDB)^2^, numerous methods have been developed based on different scientific disciplines, ideas, and various computational techniques. In the past few years, methods that use machine learning methods, particularly deep neural networks, made a large improvement in structure prediction accuracy in the Critical Assessment of techniques in protein Structure Prediction (CASP)^3^. In CASP14, a breakthrough was achieved by AlphaFold2^1^, which showed the best performance among participants with a substantial gap to the second-best method. Remarkably, the accuracy of AlphaFold2 models often reaches what would be expected from X-ray crystallography. It has been reported that models generated by AlphaFold2 have indeed helped experimental protein structure determination, as such models were successfully used for molecular replacement in X-ray crystallography and for density interpretation of cryo-EM maps^4,5^.

Soon after the release of the AlphaFold2 code, predicted structure models by AlphaFold2 for proteins from 21 major model organisms have been released at the AlphaFold Protein Structure Database^6^. This is an invaluable resource for the biology community as modelled protein structures can be easily obtained without installing and running the AlphaFold2 software. Many proteins that do not have experimentally determined structures now have computational models with an expected high accuracy.

Here, we provide the infrastructure, 3D-AF-Surfer, for real-time protein structure model search within AlphaFold2 models and across entries in PDB at https://kiharalab.org/3d-surfer/submitalphafold.php. In any database, the functionality for quick entry search and comparison are essential. In 3D-AF-Surfer, quick structure search against the entire PDB and AlphaFold2 models is realized with 3D Zernike descriptors (3DZD), which are rotationally invariant, mathematical representations of 3D shapes^7^. 3DZDs were shown to be effective in rapid protein structure database search^8^ and other tasks that involve biomolecular shape comparison and matching^9^. In 3D-AF-Surfer, we further developed neural networks that take 3DZDs of proteins as input and achieve more accurate retrieval of proteins of the same fold than direct comparison of 3DZDs.

In 3D-AF-Surfer, protein structure models generated by AlphaFold2 for 21 proteomes were retrieved from the European Bioinformatics Institute’s FTP server of the AlphaFold Database (https://ftp.ebi.ac.uk/pub/databases/alphafold) on July 22, 2021, which is still up-to-date on Oct 21, 2021. AlphaFold2 assigns one of four confidence levels, from very high confidence to very low confidence, to each amino acid position in a model. The confidence levels were assigned by predicted local distance difference test (pLDDT) score^10^, which examines the accuracy of Cα atom distances in a model. Since many models have low or very low confidence regions, which often have unfolded conformation, we extracted confident domain region(s) from each model in 3D-AF-Surfer (see Methods). In total, this procedure yielded 508,787 domains, which cover 48.8% of residues in the all the Alphafold2 models. The statistics of model counts is provided in Supplementary Table 1.

Fig. 1 illustrates the input and output panels of 3D-AF-Surfer, available at https://kiharalab.org/3d-surfer/submitalphafold.php. In the input panel, users can enter the AlphaFold model ID, PDB ID or upload the file of the query structure (Fig. 1a). When the first couple of letters of ID are entered, candidates of the rest will be listed. Then, the representation of protein structures used to compute 3DZD needs to be specified (full atom or mainchain). Next, select the database to search against, which can be the full AlphaFold proteome database, structures from PDB (complexes, domain structures) or both. Users also have an option to select the method of the database search, a deep neural network-based search (the default setting), which is suitable for retrieving proteins with the same fold (see below) or original 3DZD-based search that is equipped in 3D-Surfer. The result page shows a table where the query structure is displayed on the left side and on the right, and a list of retrieved structures ranked by their similarity to the query (Fig. 1b). Clicking a retrieved structure invokes a new search using the selected structure as the query. The panel also provides the option to compute the root mean square deviation (RMSD) between the query and the displayed similar structure. Pockets in the query structure can be identified using VisGrid^11^ or LIGSITE^12^. Finally, shown at the bottom of the page is the 3DZD of the query structure.

**Figure 1.**
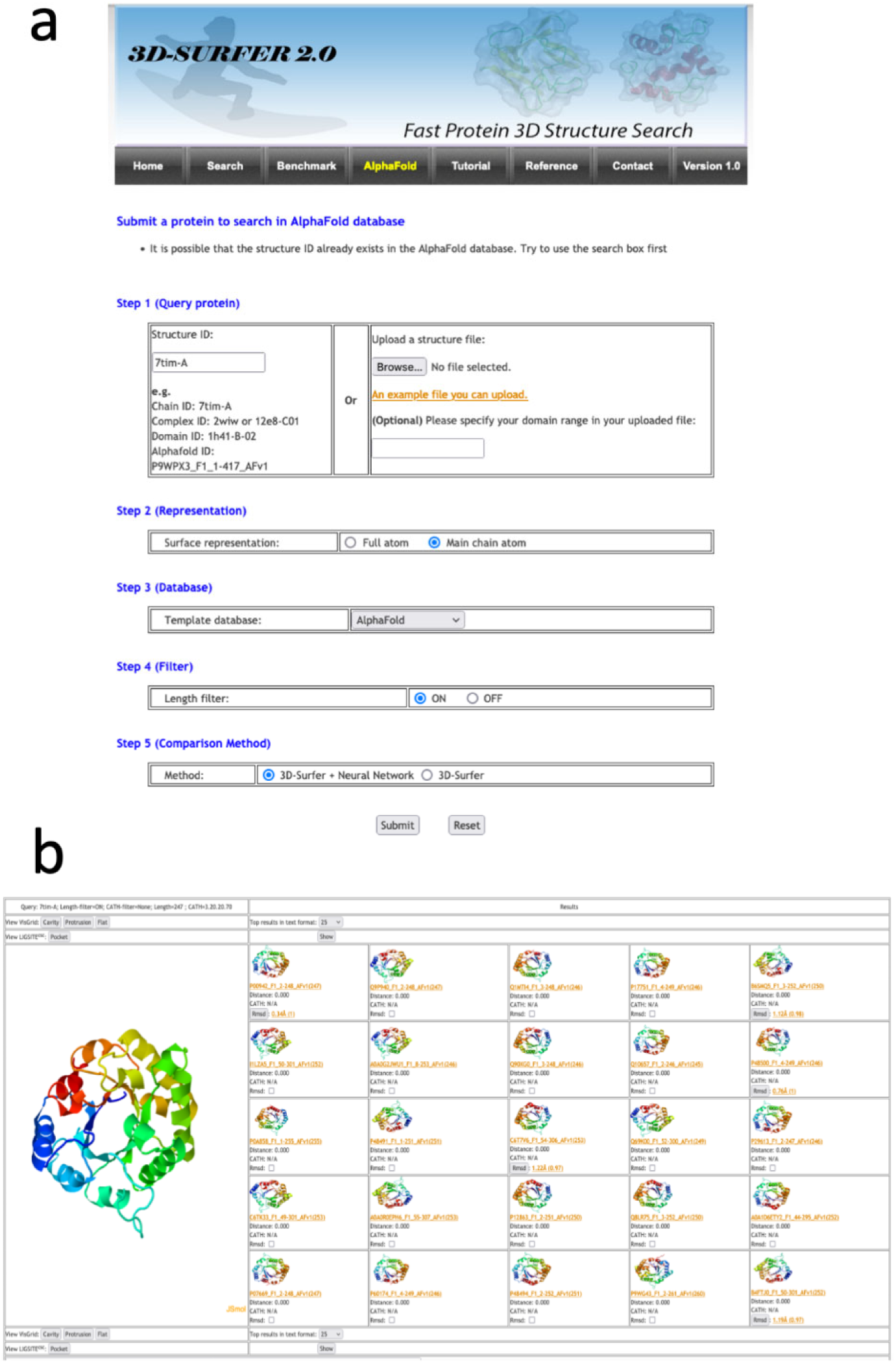
Input and an output example of 3D-AF-Surfer. **a**, the input page (see text). **b**, an example output page. The query was PDB ID: 7tim-A, a TIM-barrel fold and search was against AlphaFold models using the deep neural network. As shown, retrieved top 25 hits are all TIM-barrel folds with a distance of 0.0, indicating that the network judged that these structures are highly likely to belong to the same fold.

As of October 21, 2021, PDB entries in 3D-AF-Surfer are updated weekly. Currently, the server holds 539,936 protein chains and 249,091 additional domain structures from PDB, and 508,787 domain structures from the AlphaFold Database. Average time for a search measured over ten queries is as follows, when the neural network is used: Against AlphaFold domains: 55 seconds (s); PDB chains: 1 min 10s; PDB domains: 22s; PDB chains+domains: 1 min 15s; All of the above: 2 min 26s. Search is faster if 3DZD is used: 3s against AlphaFold domains; 1.35s, 1.45s, 1.93s against PDB chains, domains, and chains+domains, respectively, and 2.45s for All of the above.

Fig. 2a shows a breakdown of fold class of domain structures of AlphaFold2 models in comparison with SCOPe^13^. Four fold classes are considered, α, β, αβ, and small proteins. αβ corresponds to the α+ β and α/β fold classes in SCOPe. The fold classification was performed with a machine learning method, a bagged ensemble of support vector machine classifiers (SVMs) using the secondary structure content of SCOPe domains (see Methods). The bagged ensemble had an accuracy of 91.5% (Supplementary Table 2). The classification result for SCOPe (Fig. 2a) is qualitatively consistent with earlier statistics of CATH^14^, where the αβ class occupies over 50% and the share of α-class is around 15%. On the other hand, we note a greater prevalence of α-class structures among the AlphaFold2 domains (Fig. 2b) than in the SCOPe statistics (Fig. 2a). This result probably indicates that α-class structures tend to have higher confidence than other classes.

**Figure 2.**
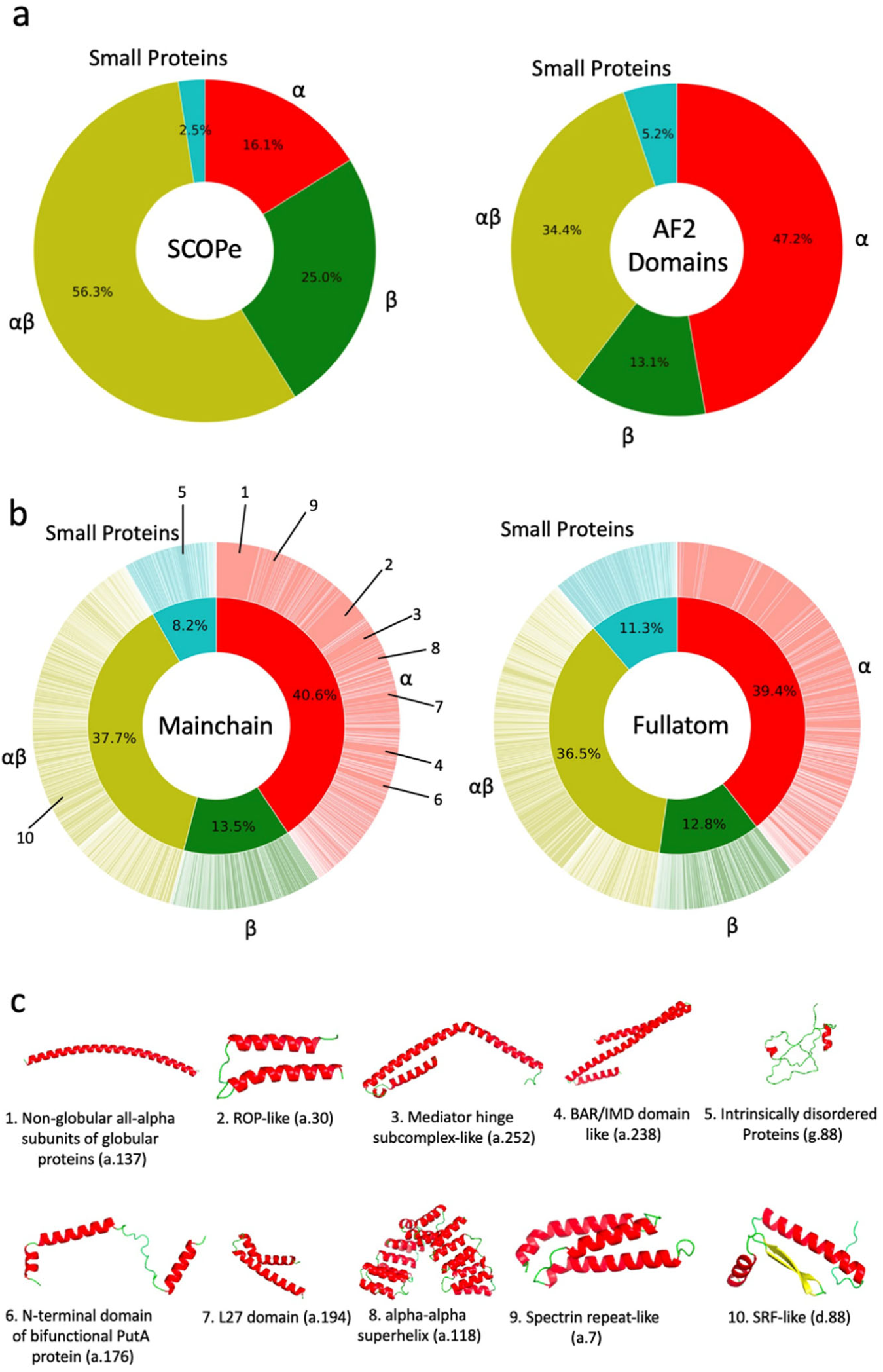
Distribution of protein secondary structure classes and fold classes of confident domains of AlphaFold2 models. **a**, the secondary structure classes were assigned to SCOPe domains and domains of high confidence in AlphaFold2 models. Four classes were considered, α, β, α β, and small proteins. Left, SCOPe (232,630 domains); right, domains of high confidence in AlphaFold2 models. (508,787 domains). The classification was performed using a bagged SVM ensemble (see Methods). SCOPe domains (left) were also classified with the SVM ensemble to be able to compare with the results on AlphaFold2 domains (right). **b**, fold classification of the AlphaFold2 structure domains of high confidence. The classification was performed with the deep neural networks that were trained on the fold assignment provided in SCOPe (see Methods). The outer wheel indicates fraction of each fold. Folds were ordered according to SCOPe IDs. Left, the fold distribution of AlphaFold2 domains using the deep network trained on 3DZDs of full atom domain structure surface. The inner wheel shows fraction of secondary structure classes. Since this classification was based on the fold assignment, the fractions are overall consistent but not identical to those shown in panel a. The top 10 most abundant folds are indicated. Right, the fold distribution using the deep network trained on 3DZDs of surface shapes with main-chain atoms. **c**, the 10 most abundant folds among AlphaFold2 domains. The fraction of each fold is indicated in the wheel diagram on the left in panel b. For each fold, an example of AlphaFold domains is shown. 1. Non-globular all-alpha subunits of globular proteins (a.137). Example shown is A0A1D6E4Z3_F1, residue 823-895 (maize). 2. ROP-like (a.30): A0A1D6MV33_F1, residue 758-815 (maize). 3. Mediator hinge subcomplex-like (a.252). Q4DL50_F1, residue 384-495 (*T. cruzi*). 4. BAR/IMD domain-like (a.238). Q8LE58_F1, residue 2-133 (*Arabidopsis*). 5. Intrinsically disordered proteins (g.88). I1L2C2_F1, residue 210-284 (soybean). 6. N-terminal domain of bifunctional PutA protein (a.176). A7MBM2_F1, residue 157-225 (human). 7. L27 domain (a.194). A0A1D6PKM6_F1, residue 314-375 (maize). 8. alpha-alpha superhelix (a.118). K7KHY8_F, residue 213-524 (soybean). 9. Spectrin repeat-like (a.7). P38637_F1, residue 149-238_AFv1 (*S. cerevisiae*). 10 SRF-like (d.88). A0A1D6NUQ9_F1, residue 2-74 (maize).

To have an overall grasp of the fold distribution of AlphaFold models, we used the deep neural network of 3D-AF-Surfer and classified AlphaFold domain structures into SCOPe folds (Fig. 2b). For this classification, we considered 1,101 folds in the class a (all α proteins), b (all β proteins), c (α/β proteins), d (α+β proteins), and g (small proteins) in the SCOPe database. The neural network takes 3DZDs of two protein structures and outputs the probability that the two structures belong to the same SCOPe fold^15^ (Supplementary Figure 1; see Methods). We trained two networks, one that uses 3DZDs computed from full-atom protein surface and another one that takes 3DZDs computed from main-chain Cα, C, and N atoms^16^. The network with the main-chain atoms showed higher classification accuracy (95.0%) than the full-atom network (Supplementary Table 3). This accuracy was higher than the original 3D-Surfer^8^, which compares 3DZDs directly with the Euclidean distance.

The fold classification results are shown in Fig. 2b. The inner and the outer wheels of the pie charts show the classification result at the secondary structure class level and at the individual SCOPe folds, respectively. The distribution of the secondary structure class levels is consistent with Fig. 2a, which was classified from secondary structure content of models. Classifications using the main-chain atoms (the left panel in Fig. 2b) and full-atoms (the right panel) were also consistent. Overall, the α-class folds are dominant when all the proteomes are considered. Among the 10 most abundant folds (Fig. 2c), eight of them belong to the α-class, one to the αβ-class (d.88), and one to the small protein class (g.88), respectively. When fold classification was examined at each organism (Supplementary Table 4), some differences among organisms were observed. Four bacterial species, *M. jannaschii, M. tuberculosis, S. aureus*, and *E. coli* have four to five αβ-class folds among top 10, which include TIM β/α-barrel (c.1) and PLP-dependent transferase-like fold (c.67). Immunoglobulin-like β-sandwich (b.1) was within top 5 in mouse, rat, and human. Plant proteomes had a β-class fold, CsrA-like (b.151) among top 10.

At last, we also analyzed low-confidence regions of AlphaFold2 models as they are not handled in 3D-AF-Surfer and thus left out from the above analysis. Particularly, we analyzed correlation between the low-confidence regions (pLDDT ≤ 0.5 and 0.7) from AlphaFold2 models and disorder predictions. We used two disorder prediction methods, SPOT-Disorder-Single^17^ and flDPnn^18^. According to the two methods, about 14% to 18% of residues are disordered (Fig. 3a). On the other hand, considering 0.5 and 0.7 pLDDT as cutoffs, more residues, 25% and 36.5%, in AlphaFold2 models were in low confidence regions (Fig. 3b). The percentage of low-confidence residues varies for different species. Low-confidence regions are relatively small (7-13%) in the four bacterial proteomes, while *D. discoideum* has the largest fraction of low-confidence residues, 58.4%. For the other organisms, low-confident residues share about 30-40%.

**Figure 3.**
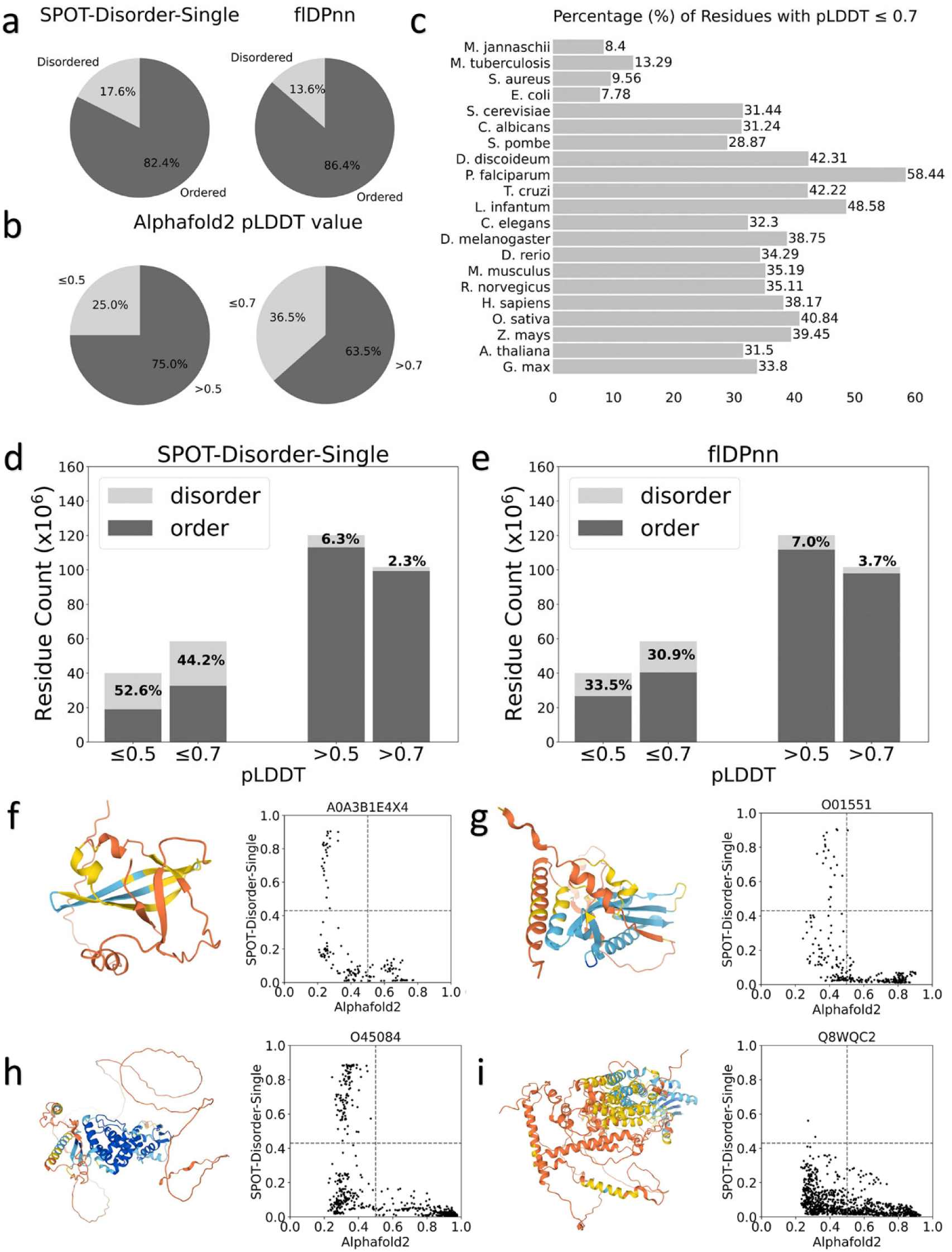
Correlation between predicted disordered regions and low-confidence regions in AlphaFold2 models. **a**, percentages of residues that were predicted as disordered or ordered by SPOT-Disorder-Single (left) and flDPnn (right). **b**, percentages of residues that were with a low confidence score ≤ 0.5 (left) and ≤ 0.7 (right). **c**, percentages of residues with a low confidence score ≤ 0.7 for each proteome. **d**, the number of residues in predicted disordered regions in low-confidence regions with 0.5, 0.7 cutoff. prediction was made by SPOT-Disorder-Single. **e**, the same type of analysis as panel d using disorder region prediction by flDPnn. **f, g, h, i**, case studies of correlation between the confidence score and disorder propensities by SPOT-Disorder-Single. The AlphaFold2 model ID is provided at the top of the plot. Left, the model structure. The color code shows the confidence level as used in the AlphaFold Database: blue (pLDDT>90), light blue (90>pLDDT>70), yellow (70>pLDDT>50), orange (pLDDT<50). Right, correlation between the confidence score (x-axis) and disorder propensity (y-axis) for each residue by SPOT-1D-Single.

In Fig. 3d and e, we compared disorder predictions and the model confidence scores using two score cutoffs, pLDDT of 0.5 and 0.7. When SPOT-Disorder-Single was used for disorder prediction (Fig. 3d), 52.6% and 44.2% of low-confidence regions defined with a pLDDT cutoff of 0.5 and 0.7, respectively, were predicted as disordered. Thus, reversely, 47.4% and 55.8% of low-confidence regions were predicted as ordered. On the other hand, almost all high confident-regions were predicted to be ordered. The result was essentially the same when flDPnn was used (Fig. 3e), except that disordered residues in low-confidence regions became even less, 33.5% and 30.9% using pLDDT of 0.5 and 0.7 as a cutoff, respectively. The results indicate that low-confidence regions do not always correspond to disordered regions, at most only 30 to 50%, and rest would be folded in native protein structures. Fig. 3f-i show several examples. The first three panels (f, g, h) are similar cases. Low-confidence residues at pLDDT around 0.4 or lower have a wide range of disorder propensities, and about half of such residues have low disorder propensity and probably would be folded in the native structures. In the model shown in Fig. 3i does not have residues with high disorder propensity, implying that the protein would be well folded in the native form.

## Online Methods

### Extraction of confident domain regions in Alphafold2 models

To extract a confident domain in an AlphaFold2 model, we first extracted all contiguous regions of more than 50 confident residues that have a pLDDT score greater than 70.0. Then, confident regions separated by at most 5 non-confident residues were merged, along with the intervening residues regardless of confidence level. AlphaFold2 models were discarded if they have no confident domains. In total, this procedure yielded 508,787 domains. 83,615 (22.9%) models out of 365,198 total AlphaFold2 models contain no confident domains. The statistics of model counts is provided in Supplementary Table 1. In terms of total residues, the domain dataset in 3D-AF-Surfer contains 48.8% (78,133,986 residues) of residues among the residues in the all the AlphaFold2 models (160,235,650 residues).

### SCOPe Benchmark Dataset for Structure Classification

We downloaded the latest version of the SCOPe dataset release 2.07 from the download page of the SCOPe website (https://scop.berkeley.edu/downloads/). The dataset included 256,391 structures in 1,430 folds after removing structures in class I (Artifacts). For each of the protein structures we used EDTSurf^19^ to generate the solvent excluded surface, for which a 3DZD vector is computed. We computed two types of 3DZD vector for a structure. The first one is computed using full atom of the protein structure. The second 3DZD is computed using only the main-chain Cα, C, and N atoms from the structure, because this main-chain surface representation performed better in our previous work^16^.

### Classification of secondary structure class with bagged SVM

The fold classification was performed with a bagged ensemble of SVMs using the secondary structure content of SCOPe domains. In bagging, *N* = 20 different classifiers were trained on 5% of the SCOPe dataset selected randomly with replacement. The output classes were then decided by voting. On the training set, the bagged ensemble had an accuracy of 91.5%. This accuracy was higher than five other methods we compared, which were a multinomial logistic regression, two SVM architectures, and two expert-designed approaches. In the expert-designed approaches, the secondary structure content thresholds, i.e. fraction of amino acids in a protein in α helices, β strands, and coil (other structures) were considered. A detailed comparison of these methods is provided in Supplementary Table 2.

### 3D Zernike Descriptors (3DZD)

3DZDs are mathematical rotation-invariant moment-based descriptors. For a protein structure, a surface from a set of atoms was constructed and then mapped to a 3D cubic grid of size N^3^ (N = 200). Each voxel (a cube defined by the grid) is assigned either 1 or 0; 1 for a surface voxel that locates closer than 1.7 grid intervals to any triangle defining the protein surface, and 0 otherwise. This grid was considered as a 3D function *f* (**x**), for which a series was computed in terms of the Zernike-Canterakis basis^7^. 3D Zernike moments of *f* (**x**) are defined as the vector of coefficients of the expansion in this orthonormal basis, and the rotationally invariant 3DZDs are defined as norms of the vectors^9^.

### Deep neural network for fold classification

Using the generated 3DZD, we trained a deep neural network that outputs the probability that a given pair of protein structures belong to the same fold. The network (Supplementary Figure 1) takes the 3DZDs of two protein shapes as input. Three hidden layers have 250, 200, and 150 neurons, respectively, which were used as the encoding of an input 3DZD. The encoder is connected to the feature extractor, a fully-connected network, which takes the 3DZDs of the two proteins, and the encodings from the three hidden layers, and four metrics that compare two vectors, the Euclidian distance, the cosine distance, the element-wise absolute difference, and the element-wise product, and the two features of the two protein shapes (the difference in the number of vertices and faces). In total, the number of the input features of the feature comparator is 2*121 + 2 * (250 + 200 + 150) + 2 * 4 + 2 = 1,452 features. The first term is the 3DZDs of order 20 (n=20), which is a 121-element vector of the two protein shapes. The third term, 2 * 4 comes from the four-comparison metrics applied to two representations of the two proteins, the original 3DZDs and encodings, which concatenate the output of the input layer and the three intermediate layers of the encoder. The feature comparator outputs a score between 0 and 1 using a sigmoid activation function, which is the probability that the two proteins are in the same fold classification in the SCOPe database.

The training and validation were performed on the aforementioned structure dataset of SCOPe. Out of 256,391 structures in 1,430 unique folds, we set aside 2,541 structures for model validation. For each of the structure in the database, we generated positive and negative pairs. Positive pairs are protein structures that belong to the same fold, while negative pairs are from different folds. For training, we randomly sampled a balanced set of positive and negative pairs based on the batch size (i.e. 32 positive pairs and 32 negative pairs for a batch size of 64). We used ADAM for parameter optimization with a binary cross entropy loss function. The learning rate was explored from 1e-3 to 7e-3 and 0.1-0.7 in our previous work and set to 0.005^15^. The accuracy of networks was evaluated on the negative and positive set generated from the 2,541 structures, which totals 167,872 pairs.

To assign a fold to a query protein, the query was compared with 10 randomly selected structures from each SCOPe fold. Then, the fold that showed the highest probability for the query is assigned. Although the training of each network was performed on the folds for all the classes except for the artifact class (class I), in the pie charts in Figure 2 we assigned to folds that belong to α, β, α β (α+β and αβ), and small proteins, because the other classes are consider factors other than structural features.

### Disorder region prediction methods

We used two methods, flDPnn^18^ and SPOT-Disorder-Single^17^. flDPnn uses profile information computed by three other methods, which is processed by a deep learning architecture to output residue-wise disorder prediction. flDPnn showed the top performance in the most recent Critical Assessment of protein Intrinsic Disorder prediction (CAID) experiment^20^. Following the instruction of the software, residues with a disorder propensity score above 0.3 were considered as disordered. We used the open-sourced implementation and trained models at http://biomine.cs.vcu.edu/servers/flDPnn/.

SPOT-Disorder-Single is a fast method that computes prediction from the single sequence of the query. It uses an ensemble of nine models. At their core, each model is constructed from ResNet blocks and/or LSTM BRNN blocks. Following the instruction of the software, residues with a disorder propensity score above 0.426 were considered as disordered. We adopted the local version of SPOT-Disorder-Single available at (http://sparks-lab.org/server/SPOT-Disorder-Single) and kept the default configuration.

## Acknowledgement

This work was partly supported by the National Institutes of Health (R01GM133840, R01GM123055, and 3R01GM133840-02S1) and the National Science Foundation (CMMI1825941, MCB1925643, and DBI2003635).

## Notes

### Competing Interest Statement

The authors have declared no competing interest.

### Summary of Updates

There was a mistake in one of the web addresses in the manuscript, and we corrected it.

